# Reverse engineering placebo analgesia

**DOI:** 10.1101/2024.02.12.579946

**Authors:** Bin Chen, Nitsan Goldstein, Julia Dziubek, Shengli Zhao, Andrew Harrahill, Akili Sundai, Seonmi Choi, Vincent Prevosto, Fan Wang

## Abstract

Placebo analgesia is a widely observed clinical phenomenon. Establishing a robust mouse model of placebo analgesia is needed for careful dissection of the underpinning circuit mechanisms. However, previous studies failed to observe consistent placebo effects in rodent models of chronic pain. We wondered whether strong placebo analgesia can be reverse engineered using general anesthesia-activated neurons in the central amygdala (CeA_GA_) that can potently suppress pain. Indeed, in both acute and chronic pain models, pairing a context with CeA_GA_-mediated pain relief produced robust context-dependent analgesia, exceeding that induced by morphine in the same paradigm. We reasoned that if the analgesic effect was dependent on reactivation of CeA_GA_ neurons by conditioned contextual cues, the analgesia would still be an active treatment, rather than a placebo effect. CeA_GA_ neurons indeed receive monosynaptic inputs from temporal lobe areas that could potentially relay contextual cues directly to CeA_GA_. However, in vivo imaging showed that CeA_GA_ neurons were *not* re-activated in the conditioned context, despite mice displaying a strong analgesic phenotype, supporting the notion that the cue-induced pain relief is true placebo analgesia. Our results show that conditioning with activation of a central pain-suppressing circuit is sufficient to engineer placebo analgesia, and that purposefully linking a context with an active treatment could be a means to harness the power of placebo for pain relief.

## MAIN TEXT

Placebo analgesia is a widely observed phenomenon. In clinical settings, placebo analgesia refers to the ability of an inert treatment to relieve pain as a result of expectations, beliefs, and/or environmental and psychosocial contexts surrounding the treatment for patients^1–3^. In randomized controlled clinical trials, placebo analgesia occurred in 10%–60% of chronic pain patients^4–6^. Harnessing the power of the placebo could therefore boost active treatment to cut medication intake, reduce severe side-effects from medications, and decrease the financial costs for patients^7,8^. To exploit the proper application of placebos in clinical practice, we need a thorough mechanistic understanding of this phenomenon. Currently, studies suggest that expectation and associative learning (classical conditioning) are two key psychological mechanisms underlying the placebo effect^2,3,9^. It is believed that administering a placebo treatment to a patient triggers a psychological response shaped by the contextual factors surrounding the treatment, such as the hospital setting and interactions with doctors. This context may evoke recollections of positive outcomes from previous treatments in a similar context, leading to positive expectations. Such expectations in turn manifest as an actual improvement in the patient’s well-being, i.e. placebo effect^10^. At the molecular level, when placebo analgesia is induced with repeated exposure to opioid drugs, the effect is blocked by the opioid antagonist naloxone, suggesting the involvement of endogenous opioids. However, when placebo is induced by repeated exposure to nonopioid agents, such as nonsteroidal anti-inflammatory drugs (NSAIDs), the effect is insensitive to naloxone blockade and instead depends on cannabinoid signaling^10,11^. The exact neural circuits mediating opioid or non-opioid placebos remain poorly defined. Human imaging studies have observed multiple brain regions either activated or deactivated during placebo analgesia^12^, although most of these studies focused on acute but not chronic pain conditions.

A robust rodent model of placebo analgesia would allow researchers to use modern molecular and genetic tools to dissect the circuit mechanisms underlying placebo analgesia. Most of the limited animal studies on placebo used repeated exposures to opiates such as morphine paired with various cues (olfactory, taste, visual or contextual) to elicit placebo analgesia (see review^13^). Interestingly, one study found a nocebo, rather than a placebo, effect induced by a cue paired with morphine^14^. Since morphine is systemically administered and opioid receptors are widely expressed, it is difficult to pinpoint the exact circuits and mechanisms that produce the placebo effect using this paradigm. Another major unsolved obstacle is that placebo analgesia in rodents has been consistently shown in acute pain models, but is highly unreliable in chronic pain models. For example, McNabb et al. did not observe any placebo analgesia in a rat model of chronic neuropathic pain using spinal nerve ligation (SNL), and Yin et al. failed to elicit placebo analgesia in a CFA-induced long-lasting inflammatory pain model^15,16^. On the other hand, Zeng et al. found that 36% of rats showed placebo responses following SNL^17^. It is therefore unclear whether it is possible to produce consistent and strong placebo effects in chronic rodent pain models which have greater translatable validity to chronic pain in humans.

Establishing a consistent circuit-based placebo analgesia model will help future dissections of detailed underlying mechanisms. Since we had recently found that general anesthesia activated neurons in the central amygdala (CeA_GA_) have potent anti-nociception effects when activated, we asked whether we can reverse engineer placebo analgesia by activating these neurons. To do this, we paired a specific context with the activation of CeA_GA_ neurons (pain relief) in acute and chronic mouse pain models, and subsequently tested whether the paired context alone was sufficient to reduce pain-related hypersensitivity. We found that pairing a context with activation CeA_GA_ neurons is indeed sufficient to produce robust placebo analgesia in both acute and chronic pain models. To our knowledge, our study is the first to use circuit-specific manipulations to produce placebo analgesia.

## Results

### Placebo analgesia induced by CeA_GA_ mediated relief of acute capsaicin pain

To achieve optogenetic activation of CeA_GA_ neurons, which express Fos in response to general anesthesia, we used an activity dependent labeling technique (CANE) that we previously published^18,19^. Briefly, after Fos^TVA^ mice had undergone 2 h of isoflurane anesthesia, we bilaterally injected a mixture of CANE-lenti-Cre (pseudotyped to specifically infect Fos-positive neurons in Fos^TVA^ mice) and Cre-dependent AAV-Flex-ChR2 or GFP to express channelrhodopsin or (CeA_GA_-ChR2) or GFP control (CeA_GA_-GFP) in CeA_GA_ neurons (Fig. S1A). We first used a two-day capsaicin injection paradigm, wherein on day 1, mice were given a capsaicin injection into the right hindpaw in the “pain induction box”. Two minutes after capsaicin injection, mice were transferred to the “pain relief box” for 10 min during which optogenetic stimulation of CeA_GA_ neurons was applied. This procedure was repeated the next day (Fig. S1B). We recorded paw licking time in both the pain induction and pain relief contexts. While time spent on paw licking increased on day 2 for both groups, the CeA_GA_-ChR2 group displayed significantly less licking behavior in the relief context during laser stimulation (Fig. S1C, S1D), demonstrating strong CeA_GA_ induced analgesia as we have previously reported^19^. We next examined mechanical and heat sensitivity using von Frey and Hargreaves tests on days 3 and 4 in the absence of any laser illumination (one test per day, randomized order, Fig. S1E) in each context to test for a placebo effect. CeA_GA_-ChR2 but not control mice had significantly higher thresholds for responding to von Frey stimuli in the pain relief box compared to thresholds in the pain induction box (Fig. S1F), suggesting that the context associated with prior pain relief alone produced analgesia. However, there was no observable placebo effect on heat sensitivity (Fig. S1G).

We wondered whether the two-day capsaicin paradigm was too short to produce a strong placebo effect for both heat and mechanical sensitivity. We therefore altered the protocol to (1) increase the total time exposed to the pain relief context and (2) introduce a second context where no pain relief is provided that is distinct from the pain induction context. In this protocol, we subjected CeA_GA_-ChR2 and control CeA_GA_-GFP mice to a 6-day capsaicin conditioning paradigm (Fig. 1A, 1B). On days 1, 3, and 5, capsaicin was injected into the right paw in the pain induction box for 2 min, and mice were subsequently transferred to a box without any laser stimulation (no relief context) for 15 min. On days 2, 4, and 6, capsaicin was injected into the left paw, and mice were transferred to a different box where laser illumination of CeA_GA_ neurons was applied for 15 min (relief context). As expected, CeA_GA_-ChR2 but not control mice showed significantly less paw licking during laser stimulation in the relief context (Fig. 1C, 1D). On days 7, 8, and 9, mice were subjected to the von Frey test, Hargreave’s heat test, or conditioned place preference (CPP) tests in the relief vs. no relief context in the absence of any laser stimulation (placebo tests, Fig.1E). Notably, CeA_GA_-ChR2 but not GFP-control mice showed significantly higher mechanical threshold to von Frey fibers and longer latency to respond to heat stimuli in the relief context compared to the no relief context (Fig. 1F, 1G). Furthermore, only CeA_GA_-ChR2 mice displayed a place preference for the relief context (Fig. 1H). We noticed that CeA_GA_-ChR2 had low mechanical thresholds in the no relief box after conditioning compared to controls (Fig. 1F). This was likely due to partial toxicity resulting from repeated optogenetic activations of ChR2-expressing CeA_GA_ neurons (data not shown). Nonetheless, these results indicate that pairing capsaicin pain relief induced by CeA_GA_ activation with a context can indeed produce context-dependent placebo analgesia and strong preference for the context.

**Figure 1.**
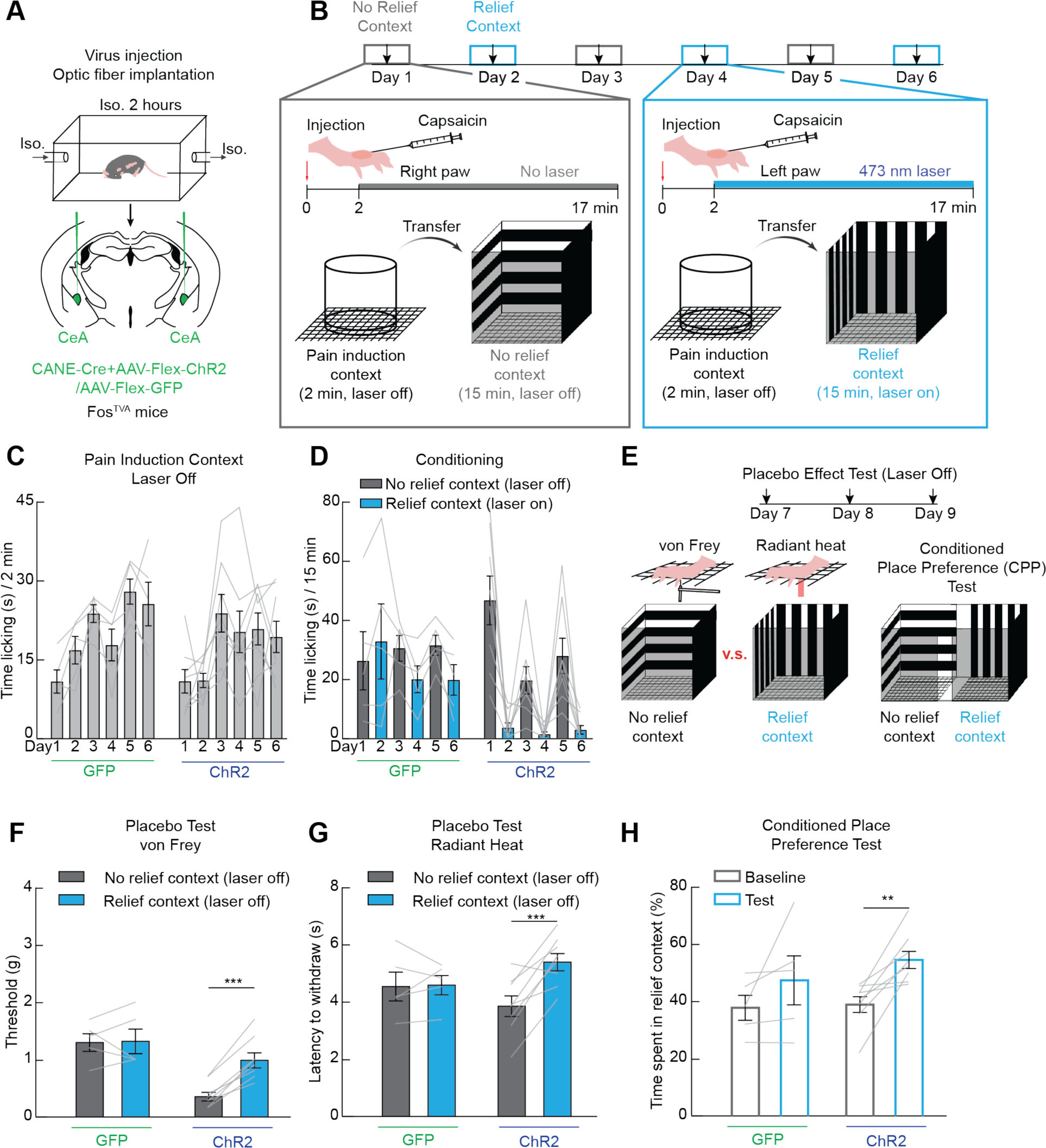
Engineering placebo analgesia with a central pain suppressing circuit in an acute capsaicin pain model. (**A**) Channelrhodopsin or control GFP was expressed in CeA_GA_ neurons by exposing mice to isoflurane anesthesia and then injecting vectors that express in recently activated neurons. (**B**) Experimental timeline used to condition animals to associate one context with CeA_GA_ activation-induced analgesia in an acute pain model (intraplantar capsaicin). (**C**) Time spent licking the injected paw in the induction chamber before laser stimulation (n=5-8/group, repeated measures two-way ANOVA, n.s.). (**D**) Time spent licking the injected paw in the conditioning chambers (n=5-8/group, repeated measures two-way ANOVA, time x group interaction p<0.001). (**E**) Experimental timeline used to test placebo analgesia by measuring mechanical (von Frey) and heat sensitivity in the no relief and relief contexts used for conditioning. Conditioned place preference (CPP) was also tested. (**F**) Withdrawal threshold during the von Frey test in GFP- and ChR2-expressing mice in the no relief context (grey) and relief context (blue, n=5-8/group, repeated measures two-way ANOVA, group x context interaction p<0.01). (**G**) Latency to withdraw from radiant heat in GFP- and ChR2-expressing mice in the no relief context (grey) and relief context (blue, n=5-8/group, repeated measures two-way ANOVA, group x context interaction p<0.05). (**H**) Time spent in the opto-stimulation-paired placebo context before and after conditioning in GFP- and ChR2-expressing mice (n=5-8/group, repeated measures two-way ANOVA, ChR2 baseline vs. test p<0.01). Data are depicted as mean ± SEM. Grey lines represent individual mice. **p<0.01, ***p<0.001.

### Benchmarking our engineered placebo against morphine induced placebo analgesia using the same capsaicin pain model

To benchmark the above placebo effect engineered with CeA_GA_ activation we examined how well morphine could induce placebo analgesia in the same acute 6-day capsaicin pain model. To do this, after establishing baseline sensory thresholds and place preference, we paired morphine injection to capsaicin treated mice with one context for 3 of the 6 days (relief context), and on alternating days, we paired saline injection with the other context (no relief context, Fig. 2A). Morphine treatment significantly reduced licking of the capsaicin-injected paw during the conditioning days (Fig. 2B) demonstrating strong analgesia. Subsequently, we tested the mechanical and heat sensitivity of these mice in the two contexts (Fig. 2C). Mice displayed decreased heat sensitivity in the morphine-paired context compared to the saline-paired context (Fig. 2D). This protocol did not consistently produce a higher mechanical threshold, nor did it enhance place preference (Fig. 2E, 2F). By contrast, our engineered CeA_GA_ mediated placebo using the same capsaicin paradigm consistently increased both mechanical and heat threshold and reliably induced place preference (Fig. 1F-H). Thus, context associated with CeA_GA_ mediated pain relief produced a more robust analgesic effect than morphine.

**Figure 2.**
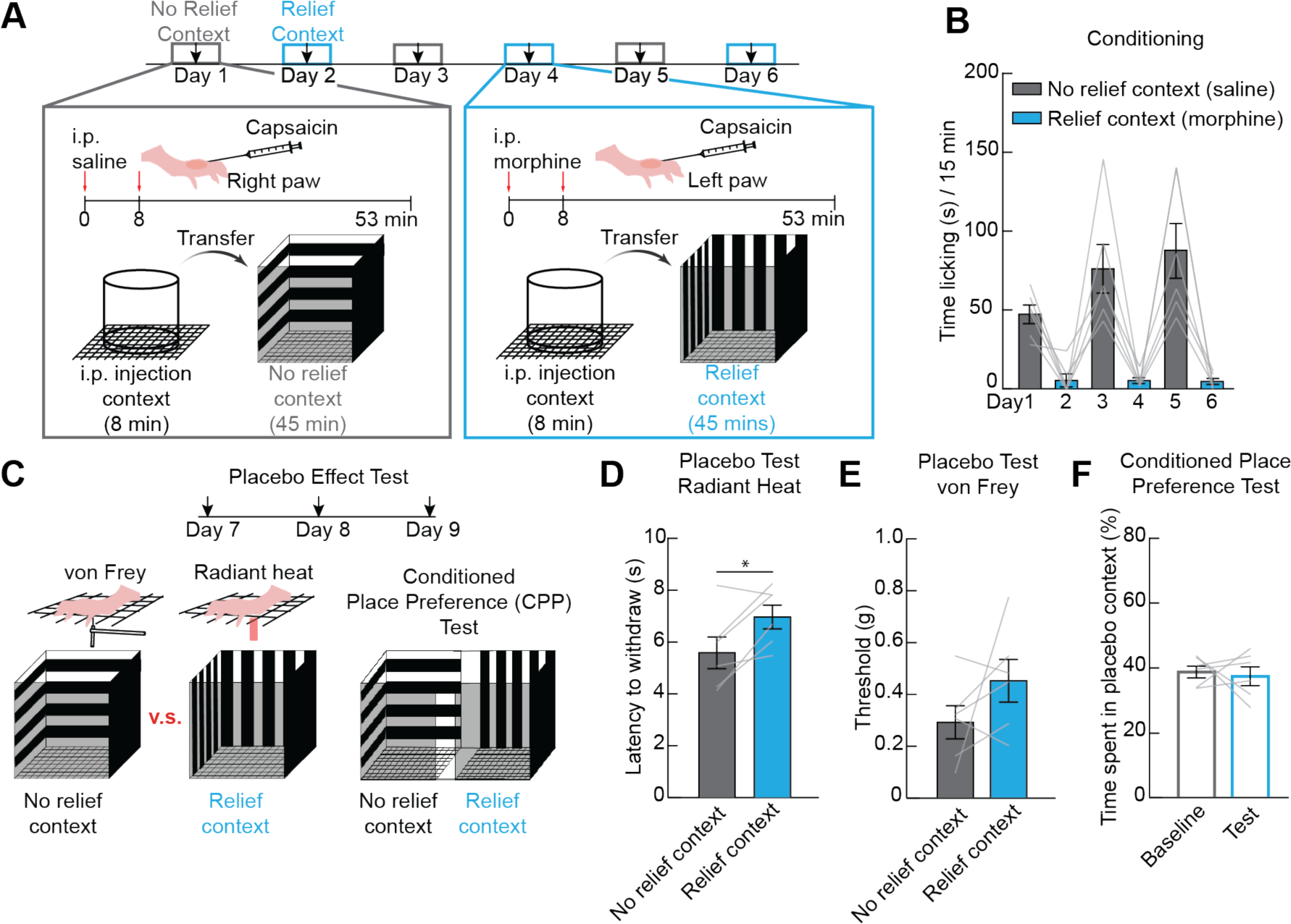
Morphine weakly induces placebo analgesia using the same acute capsaicin pain paradigm. (**A**) Experimental timeline used to condition animals to associate one context with morphine-induced analgesia in an acute pain model (intraplantar capsaicin). (**B**) Time spent licking the injected paw in the conditioning chambers (n=6, repeated measures one-way ANOVA, p<0.001). (**C**) Experimental timeline used to test placebo analgesia by measuring mechanical (von Frey) and heat sensitivity in the no relief and relief contexts used for morphine conditioning. Conditioned place preference (CPP) was also tested. (**D**) Latency to withdraw from radiant heat in the no relief saline context (grey) and the relief morphine context (blue, n=6, paired t-test, p<0.05). (**E**) Withdrawal threshold during the von Frey test in the no relief saline context (grey) and relief morphine context (blue, n=6, paired t-test, n.s.). (**H**) Time spent in the stimulation-paired placebo context before and after conditioning (n=6, paired t-test, n.s.). Data are depicted as mean ± SEM. Grey lines represent individual mice. *p<0.05.

### Placebo analgesia induced by CeA_GA_ mediated relief in a chronic pain model

We next wanted to test whether CeA_GA_ activation can induce placebo analgesia in a chronic pain model. We used the chemotherapy (paclitaxel, PTX)-induced peripheral neuropathy (CIPN) model, which is known to produce long lasting mechanical hypersensitivity. CeA_GA_-ChR2 and control CeA_GA_-GFP mice were first tested for their mechanical threshold at baseline before PTX injection. Subsequently, they were given 4 injections of 6 mg/kg PTX on days 2, 4, 6, and 8, and tested for mechanical sensitivity on day 16 (Fig. 3A). Indeed, mice became hypersensitive by day 16 as shown by their increased paw withdrawal responses to low force von Frey fibers (Fig. 3B). On days 17, 18, and 19, mice underwent conditioning training. Each morning, mice were placed in a context where laser stimulation was applied for 30 min each day (pain relief context for CeA_GA_-ChR2 mice) and in the afternoon, mice were placed in a different context with no laser application (no relief context). On day 20 we tested their mechanical sensitivity to von Frey stimuli in both contexts without laser stimulation (Fig. 3C). Remarkably, CeA_GA_-ChR2 mice experiencing chronic neuropathic pain showed strong analgesia in the relief context, where their mechanical hypersensitivity was almost completely reversed. By contrast, their hypersensitivity remained intact in the no-relief context (Fig. 3D). Control CeA_GA_-GFP mice had similar hypersensitivity in both contexts (Fig. 3D). Thus, pairing CeA_GA_ activation with a context can produce context-dependent analgesia in a long-lasting chronic pain model.

**Figure 3.**
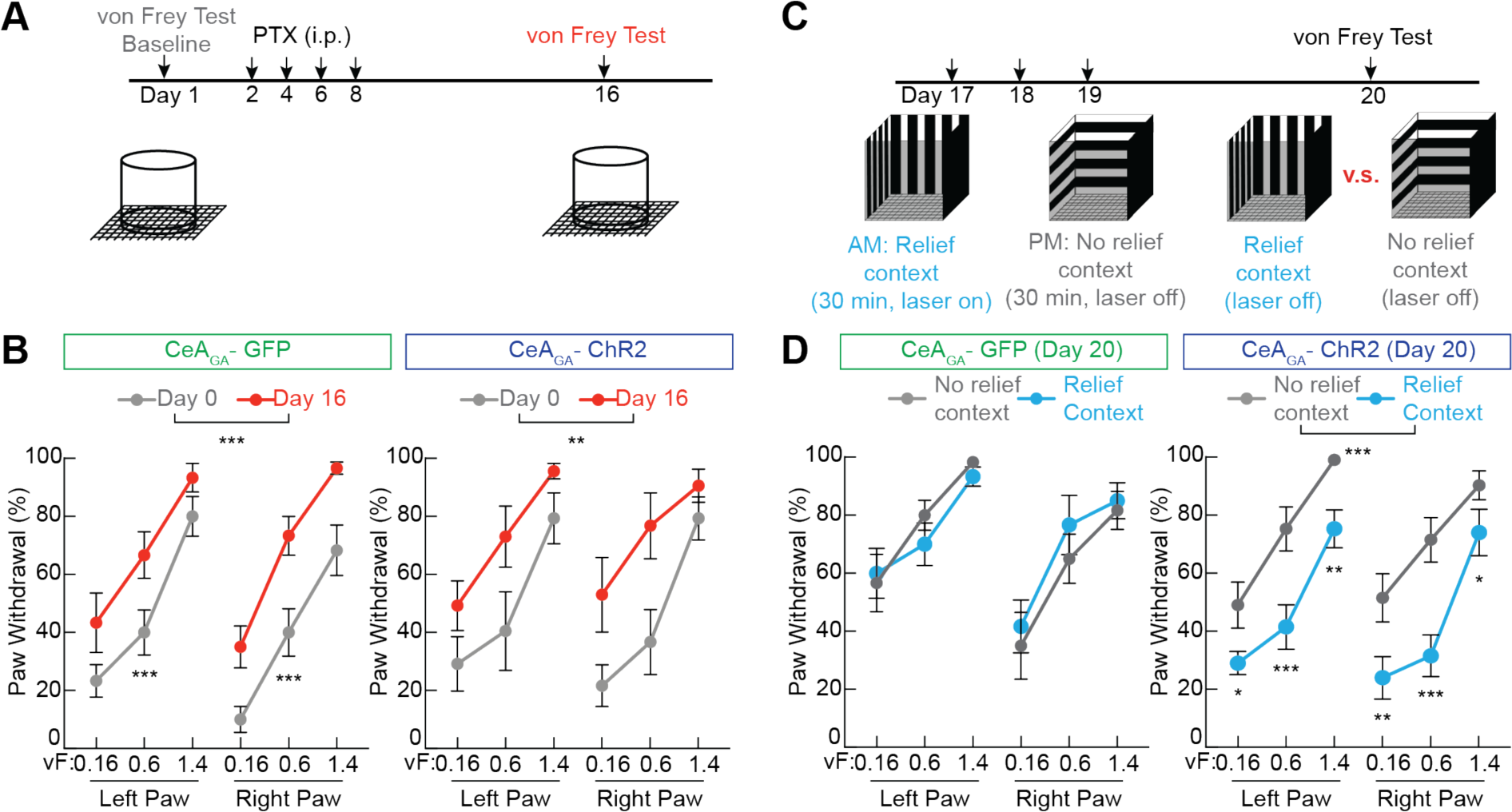
Engineering placebo analgesia with a central pain suppressing circuit in a chronic neuropathic pain model. (**A**) Experimental timeline used to induce and confirm mechanical hypersensitivity in a chronic chemotherapy-induced peripheral neuropathy (CIPN) model. (**B**) Percent of trials where a withdrawal was observed to von Frey stimuli in GFP- and ChR2-expressing mice before (grey) and after (red) treatment with the chemotherapy drug paclitaxel (n=6 (GFP), n=8 (ChR2), repeated measures two-way ANOVA, main effect of time p<0.001 (GFP), p<0.01 (ChR2)). (**C**) Experimental timeline used to condition animals to associate one context with CeA_GA_ activation-induced analgesia during CIPN and test the placebo effect with the von Frey test in each context. (**D**) Percent of trials where a withdrawal to von Frey stimuli was observed in GFP- and ChR2-expressing mice in the no relief context (grey) and relief context (blue, n=6 (GFP), n=8 (ChR2), repeated measures two-way ANOVA, main effect of context n.s. (GFP), p<0.001 (ChR2)). Data are depicted as mean ± SEM. *p<0.05, **p<0.01, ***p<0.001.

### Temporal lobe areas including hippocampus provide monosynaptic inputs to CeA_GA_ neurons

One possible simple explanation of our engineered context-dependent analgesia is that brain regions that are activated by contextual cues innervate CeA_GA_ neurons and, after associative learning, these regions enable cues to re-activate CeA_GA_ neurons to suppress pain during our testing sessions. If this were the case, the analgesia during testing sessions would still depend on an active treatment, and thus would not equate to the placebo effects observed in clinical settings. To test this possibility, we first mapped the presynaptic inputs to CeA_GA_ using monosynaptic rabies virus to determine whether any of the input regions could potentially relay contextual information to CeA_GA_. Briefly, after 2 h of isoflurane anesthesia, we co-injected CANE-Lenti-Cre with the two Cre-dependent helper viruses in CeA to express TVA-mCherry and oG (rabies glycoprotein) in CeA_GA_ neurons (Fig. 4A). Two weeks later, we injected G-deleted pseudotyped EnvA^M21^-RV-GFP (also called CANE-RV-GFP) in CeA and perfused mice 5 days after rabies injection. We observed reproducible patterns of transsynaptic tracing results (Fig. 4B-F, n=3). CeA_GA_ received wide-spread inputs from thalamic and hypothalamic nuclei, hippocampus, temporal associative cortex (TEA), and many amygdala nuclei including local inputs within CeA (Fig. 4E, 4F). Many of these regions such as hippocampus, TEA, and basolateral amygdala could potentially convey contextual information to CeA_GA_ neurons to facilitate the associative learning of context with CeA_GA_ activation and subsequent context-mediated reactivation to elicit placebo analgesia.

**Figure 4.**
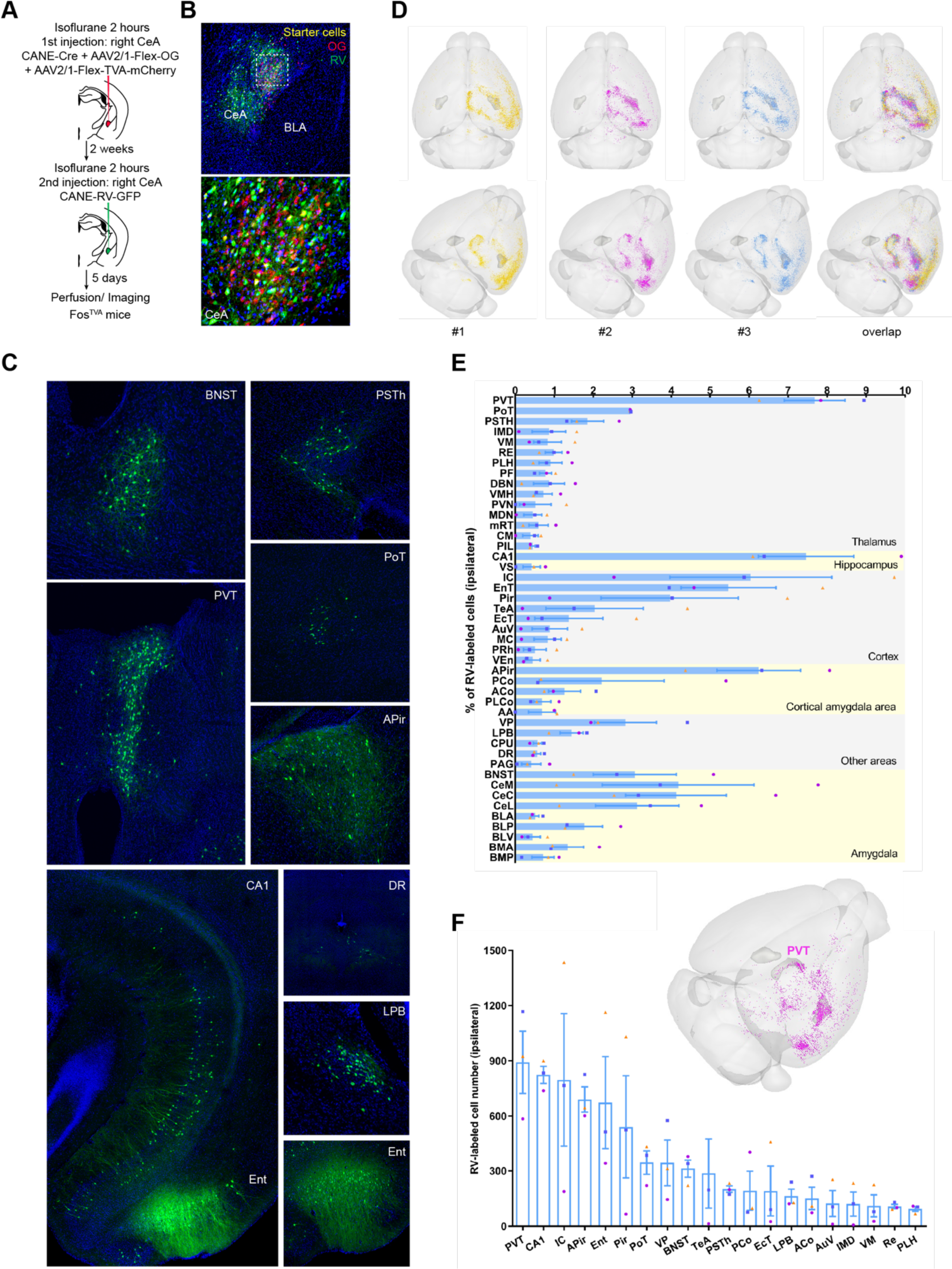
Monosynaptic inputs to CeA_GA_ neurons. (**A**) CANE targeting was used in combination with monosynaptic rabies tracing to label inputs to CeA_GA_ neurons. (**B**) Representative images of the CeA after rabies tracing. Red cells are CeA_GA_ neurons that expressed AAV helper viruses (TVA-mCherry and oG), yellow cells are starter cells that expressed both the helper and rabies viruses, and green neurons are labeled presynaptic inputs onto starter cells. The bottom image is a higher magnification view of the dotted rectangle in the top image. (**C**) Representative images showing rabies-labeled inputs from the bed nucleus of the stria terminalis (BNST), the parasubthalamic nuecleus (PSTh), the paraventricular nucleus of the thalamus (PVT), the posterior thalamic region (PoT), the amygdala-piriform transition cortex (Apir), the CA1 region of the hippocampus, the dorsal raphe nucleus (DR), the lateral parabrachial nucleus (LPB) and the entorhinal cortex (Ent). (**D**) Schematics showing the location of labeled inputs in three individual mice and the overlap among all three mice. (**E**) The percent of all rabies-labeled inputs in each brain region. (**F**) Number of rabies-labeled inputs in the most densely labeled regions. Data are depicted as mean ± SEM. Colored dots represent individual mice.

### The engineered placebo analgesia does *not* involve strong CeA_GA_ reactivation

Given the possibility of direct contextual inputs to CeA_GA_ neurons, we tested whether the placebo effect involves context-elicited re-activation of CeA_GA_ neurons. To answer this question, we needed the ability to optogenetically activate CeA_GA_ neurons during conditioning and to subsequently measure their activity during the placebo test. We therefore used CANE-Lenti-Cre in combination with AAV-DiO-jGCaMP8s-P2A-ChrimsonR-ST^20^ to express both the red-shifted opsin ChrimsonR and the green calcium indicator jGCaMP8s in CeA_GA_ neurons (Fig. 5A). This vector allowed us to stimulate CeA_GA_ with a far-red laser while image CeA_GA_ with a green laser that does not activate ChrimsonR. We confirmed the ability of ChrimsonR to activate CeA_GA_ cells with acute slice recordings (Fig. S2). These CeA_GA_-jGCaMP8s/ChrimsonR mice were then subjected to repeated PTX injections to induce CIPN and chronic pain. Afterwards, they were tested for place preference by exploring the two chambers (for 3 different sessions on two different days) while we performed fiber photometry (FP) imaging of CeA_GA_ population calcium activity. Mice were then placed in their preferred chamber in the absence of any laser stimulation for 30 min (no relief context), and then placed in their non-preferred chamber to receive ChrimsonR-mediated activation (633nm laser) for 30min (relief context). Following conditioning, we tested and confirmed placebo analgesia in the chamber paired with CeA_GA_ activation. To examine CeA_GA_ activity after conditioning in different contexts, we again let the mice freely explore the two chambers across 3 sessions while using FP to image CeA_GA_ neurons (Fig. 5A).

**Figure 5.**
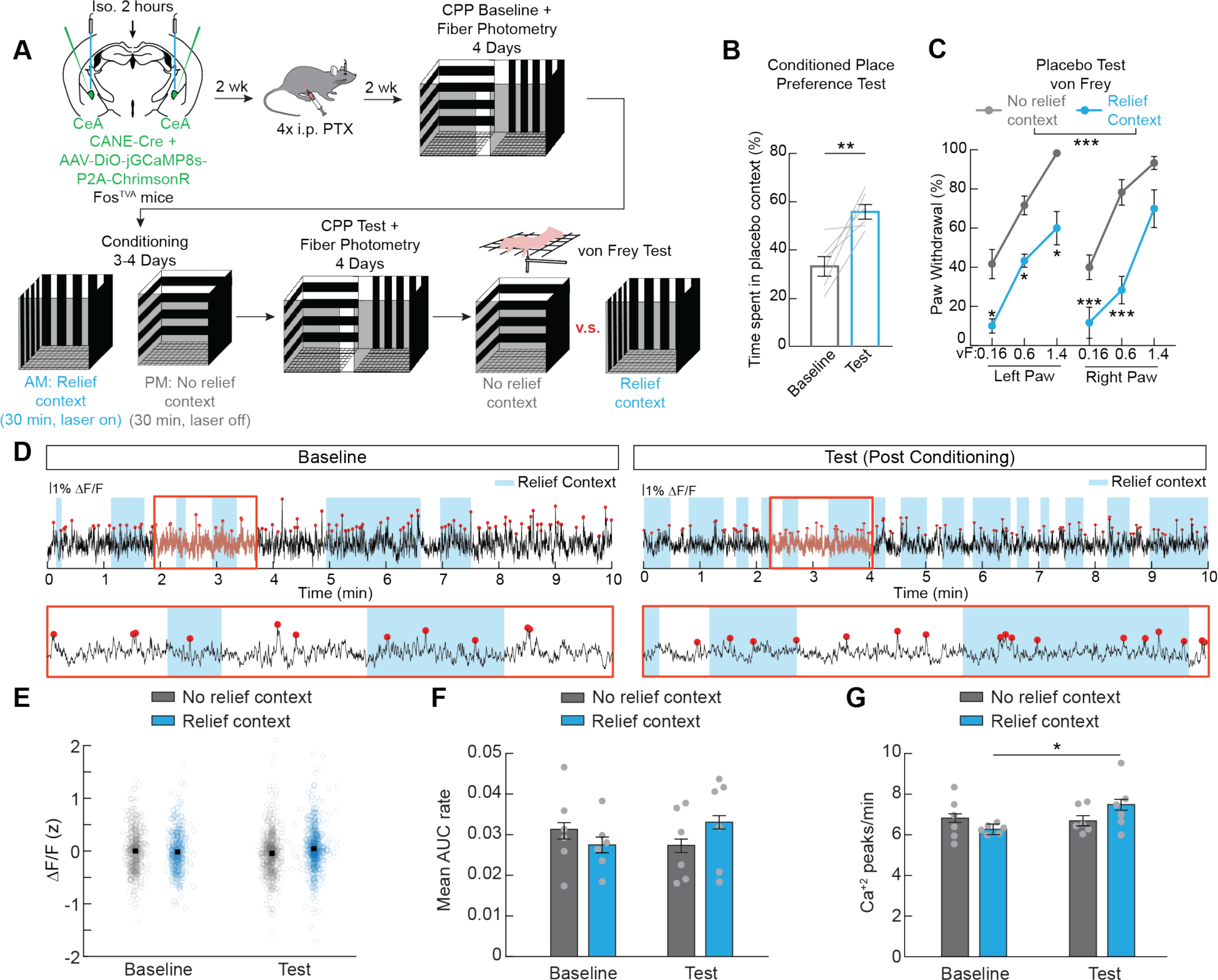
The engineered placebo analgesia does not involve strong reactivation of CeA_GA_ neurons. (**A**) Experimental paradigm to express both jGCaMP8s and ChrimsonR to induce placebo analgesia and to record activity of CeA_GA_ neurons before and after conditioning. (**B**) Time spent in the opto-stimulation-paired placebo context before and after conditioning (n=6, paired t-test, p<0.01). (**C**) Percent of trials where a withdrawal to von Frey stimuli was observed in the no relief context (grey) and relief context (blue, n=6, repeated measures two-way ANOVA, main effect of context p<0.001). (**D**) Representative GCaMP8s signal in one mouse before conditioning (baseline) and after conditioning. Blue regions show when the mouse was in the context paired with CeA_GA_ neuron activation (non-preferred chamber before conditioning, relief context after conditioning). Red dots depict statistically significant calcium events. (**E**) Average z-scored GCaMP8s signal when mice were in the no relief context (preferred before conditioning) and relief context (non-preferred before conditioning) before and after conditioning (n=6 mice, multi-factor ANOVA followed by Bonferroni post-hoc test, n.s.) (**F**) Amplitude of calcium event (area under the curve) when mice were in the no relief and relief contexts before and after conditioning (n=6 mice, multi-factor ANOVA followed by Bonferroni post-hoc test, n.s.) (**G**) Average calcium event rate when mice were in the no relief and relief contexts before and after conditioning (n=6 mice, multi-factor ANOVA followed by Bonferroni post-hoc test, p = 0.0066). Bouts where mice were in the contexts for less than 30 continuous s were excluded. Dots represent individual bouts across all mice. Data are depicted as mean ± SEM. Grey lines and dots represent individual mice. *p<0.05, **p<0.01, ***p<0.001.

We confirmed that CeA_GA_-jGCaMP8s/ChrimsonR CIPN mice established a place preference for the relief context and experienced placebo analgesia. Mice spent more time in the CeA_GA_-stimulation-paired context after conditioning during the two-chamber exploration (Fig. 5B) and their withdrawal rates during mechanical stimulation were reduced in the relief context (Fig. 5C). We then compared CeA_GA_ population FP imaging results in the relief vs. no-relief contexts before and after conditioning (Fig. 5D, Fig. S3, S4). Since the jGCaMP8s/ChrimsonR bi-cistronic vector generated low levels of jGCaMP8s expression, the resulting FP signal was noisy. We computed a 95% confidence interval for each trace, and only the calcium peaks that crossed that threshold were considered in our analysis (Fig. S3, see methods). We found that conditioning had no observable effect on the amplitude of the average fluorescence signals in the relief context as measured using either z-scored ΔF/F or area under the curve (AUC, Fig. 5E, 5F). Furthermore, there was no significant change in the calcium event rate in individual mice despite them spending more time in the relief context after conditioning (Fig. S4), although a mild increase of the calcium event rate in the pain-relief context (+1.2 events per min) but not in the no-relief context could be detected at the population level (Fig. 5G). Taken together, while there might be a slight increase in the firing probability across populations of mice, CeA_GA_ neurons were by and large not strongly re-activated in the placebo context after conditioning. These findings indicate that our reverse engineered analgesia does not depend on re-activation of CeA_GA_ mediated pain relief, hence it is a true placebo analgesia.

## Discussion

In this study, we engineered an effective contextual placebo analgesia based on pairing a context with CeA_GA_-activation mediated pain relief. In the acute capsaicin model, the analgesic effect of placebo context is generalized to both mechanical (von Frey) and heat stimuli (which were not used to establish the association between context and pain relief). In the CIPN chronic neuropathic pain model, the placebo analgesia effectively reversed the pathological mechanical hypersensitivity to the baseline observed prior to PTX treatment. In both capsaicin and the CIPN models, mice showed preference for the placebo context, indicating clear association of the context with positive affect. To our knowledge, this is the first time that a robust rodent placebo effect is established using specific circuit manipulations. Previous rodent placebo models were mostly based on opioid administration and were inconsistent in chronic pain models, while our engineered placebo is not only superior to morphine in our experimental setting, but also effective in the chronic CIPN model. This is exciting since chronic pain models like CIPN are more relevant to human pain conditions. Our study laid the foundation for future detailed dissections of cellular and circuit mechanisms underlying the generation and expression of placebo analgesia.

We further show that while CeA_GA_-activation is used to establish the placebo effect linked to specific context, the actual expression of placebo does not involve strong context-mediated reactivation of CeA_GA_. It is therefore likely that the analgesia we observed is due to the expectation of relief rather than an active treatment, suggesting that we indeed engineered a bona fide placebo analgesia. This finding also has implications for clinical purposes. For example, if an active treatment that has severe side effects can be paired with an engaging context (like an interactive app) to induce placebo, then context alone without the active treatment could be sufficient to provide relief without side effects. At present, we do not yet know how the cue-analgesia associative memory drives the placebo effect. The “remembered wellness”^21^ could involve plastic changes in distributed networks across many areas in the central and peripheral nervous systems. For example, a previous study using fMRI revealed that nociceptive processing signals in the spinal dorsal horn were reduced during placebo analgesia^22^. Future work with neural recording or imaging can help address whether, in our engineered paradigm, spinal nociceptive processing is also reduced.

We note that there are limitations and caveats in our experiments and interpretations. CeA_GA_ neurons are a heterogenous population^19^ and our fiber photometry signals are noisy. It is possible that a small subset of CeA_GA_ cells did reactivate and could account for the observed slight increase of activity at the population level and that we failed to consistently detect this with bulk population imaging. Cellular resolution imaging or opto-tagged recording will be needed in the future to resolve this possibility. In conclusion, we show that robust placebo analgesia during acute and chronic pain can be engineered by pairing a context with activation of the CeA_GA_ endogenous analgesic circuit. The expression of placebo analgesia in this paradigm is not reliant on strong re-activation of CeA_GA_, suggesting that after conditioning, context can engage circuits independent of CeA_GA_ for pain relief.

## ACKNOWLEDGEMENTS

We thank the Wang lab for helpful discussion and feedback throughout the project. This research was funded by the NIH (DE029342 to F.W.), the Yang-Tan Collectives at MIT (F.W.), and the Jane Coffin Childs Fund for Medical Research (N.G.).

## AUTHOR CONTRIBUTIONS

B.C. and F.W. initiated the project. B.C., N.G., J.D., S.Z., A.H., A.S., S.C., and V.P. performed experiments, analyzed data, and/or provided research support. B.C., N.G., and V.P. made the figures. B.C., N.G. and F.W. wrote the manuscript with comments from all authors.

## STAR* METHODS

### RESOURCE AVAILABILITY

#### Lead Contact

Further information and requests for resources and reagents should be directed to and will be fulfilled by Lead Contact, Fan Wang.

#### Materials Availability

This study did not generate new unique reagents.

#### Data and code availability

All data reported in this paper will be shared by the lead contact upon request. This paper does not report any original code. Any additional information required to reanalyze the data reported in this paper is available from the lead contact upon request.

### EXPERIMENTAL MODEL DETAILS

Adult male and female Fos^TVA^ mice (Jackson Laboratory, stock 027831^18^) were used for all behavioral experiments. Mice were housed in the vivarium with a 12-hour light and dark cycle and were given food and water ad libitum. All experiments were conducted according to protocols approved by the MIT Animal Care and Use Committee.

### METHOD DETAILS

#### Virus and reagents

CANE-LV-Cre (CANE-LV envelope [Addgene Plasmid #86666]) were produced as previously described^18^. Various AAVs were co-injected with CANE-LV-Cre: AAV2/1-hSyn-DIO-EGFP (Addgene 50457), AAV1-Ef1a-DiO-hChR2(H134R)-EYFP (Addgene 20298); AAV2/1-Flex-TVA-mCherry (Boston Children’s Hospital viral core), AAV2/1-CAG-Flex-oG (Boston Children’s Hospital viral core), EnvAM21-RV-GFP (also called CANE-RV-GFP^18^); AAV2/1-hSyn-DIO-jGCaMP8s-P2A-ChrimsonR-ST (Addgene Plasmid #174007).

#### Surgical procedures

##### Viral delivery and optic fiber implantation

To capture and express desired transgenes in CeA_GA_ neurons, Fos^TVA^ mice were anesthetized with isoflurane (1.5% isoflurane, 0.75% oxygen) for two hours (to induce Fos expression in CeA_GA_) in a chamber before mice were transported to a stereotaxic frame (David Kopf Instruments) and small craniotomies were created over the target region. The coordinates of CeA used relative to bregma were: AP = 1.20 ± 0.05 mm, ML = ± (2.86 ± 0.02) mm, DV = −4.17 or −4.22 ± 0.03 mm. The CANE-LV-Cre and Cre-dependent AAV were mixed (1:1) before injection. 0.8-1.2 µl total was delivered at 60 nl/min per injection and left for 10 minutes post-injection for efficient virus diffusion. After bilaterally viral injection, optical fibers (200 µm core diameter, RWD) were inserted 300 µm above the injection sites and secured using Metabond (Parkell) and dental cement.

##### Retrograde trans-synaptic tracing from CeA_GA_

After 2 hours of isoflurane anesthesia, we co-injected CANE-LV-Cre with the two Cre-dependent helper viruses AAV1-EF1a-FLEX-TVA-mCherry and AAV1-CAG-FLEX-oG into the CeA to express TVA-mCherry and oG in CeA_GA_ neurons. Two weeks later, mice were re-exposed to the isoflurane for 2 hours, and G-deleted pseudotyped EnvAM21-RV-GFP (also called CANE-RV-GFP) was injected into the same location. 5 days later, mice were perfused, and brains were collected.

#### In vivo optogenetic activation

Animals with optical fiber implants were connected to a 1 to 2 split branching fiber-optic patch cord (Doric) coupled to either a 473 nm or 633 nm laser (Cobolt). The light pulse was controlled by a pulse generator (Doric). The laser was applied in pulsed mode (∼3.5 mW/mm^2^, 20 Hz, 5 sec on-off) to animals that expressed ChR2 or ChrimsonR or GFP in CeA_GA_. During post hoc immunohistochemistry, viral expression, sites of injections, and insertion of optical fibers were confirmed; animals with failed expression or off-target optical fiber placement were excluded from all analyses.

#### Placebo effect engineered with CeA_GA_ activation in acute capsaicin pain model

##### Two-day capsaicin injection paradigm (Fig. S1)

2 days before conditioning, mice were habituated with handling for 5 min. On conditioning days 1 and day 2, mice were injected with capsaicin (2 µg/10 µl, surface skin of right hind paw) remained in the transparent cylinder (***pain induction context***, 6×6 inches) for 2 min, and were then transferred to the black box (***pain relief context***, 8×8×8 inches) with 473 nm laser turned on for 10 min (20 Hz, 10 ms duration, 5 sec on 5 sec off). The behavior was simultaneously recorded, and the time spent licking the injected paw in each box was manually counted. On days 3 and day 4, mice were randomly placed in either context by an experimenter. Then the other experimenter, who was blind to the group of mice, tested the pain threshold using von Frey filaments (day 3) and heat (day 4). Mice were tested in one context, and then after 2 hours, tested in the other context. The behavior boxes were positioned directly on the mesh stand without a top or bottom cover (IITC Life Science).

##### Six-day capsaicin injection paradigm (Fig. 1)

2 days before conditioning, mice were habituated to 8×8×8 inches, 2-chamber box with a separator in the middle. Mice spent 15 min in each chamber, followed by an additional 10 min of free exploration in the 2-chamber box without the separator. Each chamber contained two distinct visual patterns of horizontal and vertical stripes. 1 day before conditioning, the duration of free exploration by mice in each box was recorded in a 15 min session (Anymaze). The box that **was less preferred** by each mouse was used as the ***pain relief context***. On conditioning days 1, 3, and 5, mice were injected with capsaicin (2 µg/10 µl, right hind paw) and remained in the transparent cylinder (***pain induction context***) for 2 min; then transferred to the box (***No relief context***) for 15 min. On conditioning days 2, 4, and 6, mice were injected with capsaicin on the left hind paw and remained in the ***pain induction context*** for 2 min; then transferred to the box (***Pain relief context***) with 473 nm laser (20 Hz, 10 ms duration, 5 sec on-5 sec off) turned on for 15 min. The behavior was simultaneously video recorded, and the time spent licking the injected paw in each box was manually counted. On days 7 and 8, mice were randomly placed in either ***No relief or Pain relief context*** by an experimenter. Then, a second experimenter blinded to experimental group tested the pain threshold using von Frey filaments (day 7) and heat (day 8). Mice were tested in one context, and then after 2 hours, tested in the other context. On day 9, the duration of free exploration by mice in each context was recorded in a 15 min session (Anymaze), and the conditioned place preference was compared to the baseline.

#### Placebo effect engineered with morphine in acute capsaicin pain model

Three days before conditioning, mice were habituated to a 2-chamber box with a separator in the middle. Mice spent 15 min in each chamber, followed by an additional 10 min of free exploration in the 2-chamber box without the separator. Each chamber contained two distinct visual patterns and shapes, featuring a rectangular chamber measuring 6W×6H×8L inches with 3D gel-like smooth white wallpaper, a parallel trapezoid chamber measuring 4W×8W×8L×8H inches with black acrylic sheets. This 2-chamber box was also employed in subsequent chronic pain odel experiments. 2 days before conditioning, the duration of free exploration by mice in each box was recorded for 4 sessions (20 min each), in the morning and afternoon for 2 days. The box that **was less preferred** by each mouse was used as the ***pain relief context***. Mice were injected with capsaicin on one hind paw and remained in the ***pain induction context*** for 2 min. Following this, on 3 out of the 6 conditioning days, mice were treated with morphine (1.5 mg/ml, 15 mg/kg, i.p.), and placed in the morphine-paired box (***Pain relief context***) for a total 45 min. On alternating days, we paired saline injection with the other context (***No relief context***). The behavior was simultaneously recorded, and the time spent licking the injected paw in each box was manually counted. On days 7 and 8, mice were randomly placed in either No relief or Pain relief contexts and their pain thresholds were tested using von Frey filaments (day 7) and heat (day 8). Mice were tested in one context, and then after 2 hours, tested in the other context. On days 9 and 10, the duration of free exploration by mice in each context was recorded for 4 sessions (20 min each) and compared to baseline.

#### Placebo effect engineered with CeA_GA_ activation in a chronic pain model

We used the chemotherapy (paclitaxel, PTX, Sigma)-induced peripheral neuropathy (CIPN) model, which is known to produce long-lasting mechanical hypersensitivity. Mice were first tested for their mechanical threshold to von Frey filaments at baseline before PTX injection (day 1). Subsequently, they were given 4 injections of 6 mg/kg PTX (i.p.) every other day (days 2, 4, 6, and 8) and tested for mechanical sensitivity on day 16. On days 17, 18, and 19, mice underwent conditioning training. Each morning (10 AM-12 PM), mice were placed in a context where 473 nm laser stimulation was applied for 30 min each day (***pain relief context*** for CeA_GA_-ChR2 mice), and in the afternoon (4-6 PM, at least 5 h interval between 2 sessions), mice were placed in a different context with no laser stimulation (***no relief context***). On day 20, mice were randomly placed in either context by an experimenter. Then, the other experimenter, who was blind to the group of mice, tested the mechanical sensitivity to von Frey filaments. Mice were tested in one context, and then after 5 h, tested in the other context.

#### Fiber photometry imaging

We used CANE-LV-Cre in combination with AAV-DiO-jGCaMP8s-P2A-ChrimsonR-ST^20^ to express both the red-shifted opsin ChrimsonR and the green calcium indicator jGCaMP8s in CeA_GA_ neurons. We confirmed the ability of ChrimsonR to activate CeA_GA_ cells with acute slice recording (Fig. S2). For **fiber photometry imaging of calcium activity in CeA_GA_,** mice with optical fiber implants were connected to a low-autofluorescence 1 to 2 split branching fiber-optic patch cord (Doric) coupled to the multichannel fiber photometry (R820, RWD). 470 nm excited light and 410 nm reference light were delivered alternatively (60 Hz, 14.04 ms, 0.2 mW) during the recording with synchronized video (30 Hz, Basler). Light power was titrated based on pilot slice and behavioral experiments, to prevent ChrimsonR activation during imaging. In vivo Ca^2+^ signals of the CeA_GA_ were first recorded during exposure to isoflurane induced anesthesia (5 min baseline followed by 20 min of 1.5% isoflurane mixed with oxygen). Animals with increased Ca^2+^ signals induced by isoflurane were then subjected to repeated PTX injections to induce CIPN. Afterward, they were tested for place preference by exploring the two chambers (for 3 different sessions on two different days) while we performed fiber photometry (FP) imaging of CeA_GA_ population activity. The box that was less preferred by each mouse was used for pain relief context. In the next 3-4 days, mice were placed in their preferred chamber in the absence of any laser stimulation for 30 min (no relief context) and then placed in their non-preferred chamber to receive ChrimsonR-mediated activation (633 nm laser) for 30 min (relief context). Following conditioning, to examine CeA_GA_ activity in different contexts, the mice freely explored the two chambers (for 3 different sessions) while using FP to image CeA_GA_ and record the durations they spent in each chamber. To confirm placebo analgesia in the chamber paired with CeA_GA_ activation, we tested the mechanical sensitivity to von Frey in each context.

#### Drugs

##### Capsaicin

10 µl of capsaicin (2 µg/10 µl, Sigma-Aldrich, M2028, dissolved in normal saline with 4% ethanol and 4% Tween-80) was subcutaneously injected into the hind paw.

##### Morphine

Morphine (Sigma, M8777, 15 mg/kg) was dissolved in normal saline and injected intraperitoneally (i.p.).

##### Paclitaxel (PTX)

CIPN was induced using PTX (Sigma Aldrich T7191, 6mg/kg) given via intraperitoneal injection (i.p.) every other day for one week (4 times). 25 mg PTX was first dissolved in 700 µL DMSO, then transferred into the 3.46 mL of 50/50 Kolliphor EL (Sigma Aldrich) / ethanol solution, and then further dissolved in 0.9% saline for injections.

#### Histology

Animals were anesthetized with isoflurane, then transcardially perfused with cold phosphate buffer saline (PBS, pH 7.4), followed by 4% cold paraformaldehyde (PFA) fixation solution. All brains were post-fixed in PFA overnight at 4°C, cryoprotected in 30% sucrose PBS solution for 2–3 days at 4°C, frozen in O.C.T compound (Tissue-Tek, Sakura), and then stored at −80°C until sectioning. 80 µm free-floating coronal sections were made using a cryostat (Leica Biosystems Inc). The sections were briefly washed in PBS and stained with Dapi (1:10000, Sigma D9564) in 0.3% Triton-X100/PBS overnight at 4°C. The sections were briefly washed and mounted on slide glasses with Mowiol.

#### Image acquisition and quantification

Entire brain slices were imaged at 5x resolution with a laser scanning confocal microscope (Zeiss 700), while the CeA or interested brain regions were imaged at 10x resolution. To quantify the presynaptic neurons of CeA_GA_ neurons, the images of entire brain sections were adjusted to the coordinates of the Allen Brain atlas by using a modified custom-written code^23^. The numbers of GFP-labeled neurons in each brain section were manually counted and automatically assigned to each brain area, then normalized by the sum of total GFP-positive cells.

#### Slice Electrophysiology

##### Slice Preparation

Following isoflurane-induced anesthesia and decapitation, mice brains were swiftly extracted in partially frozen sucrose-containing artificial cerebrospinal fluid (sACSF) with the following concentrations (in mM): 90 sucrose, 60 sodium chloride, 26.5 sodium bicarbonate, 2.75 potassium chloride, 1.25 sodium phosphate, 9 glucose, 1.1 calcium chloride, and 4.1 magnesium chloride, maintaining an osmolality range of 295 to 300. Brains were then mounted and submerged in ice-cold sACSF and 300 μm-thick slices were cut using a vibratome (Leica VT1200S). Post-cutting, the slices were transferred into ACSF with the following concentrations (in mM): 125 sodium chloride, 25 sodium bicarbonate, 3 potassium chloride, 1.25 sodium phosphate, 19.5 glucose, 1.2 calcium chloride, 1.2 magnesium chloride, 1 ascorbate, and 3 sodium pyruvate, ensuring an osmolality of 300-305. The slices underwent a 45-minute recovery period in ACSF at 36 °C and were then maintained at room temperature. Both solutions were at all times saturated with carbogen (5% CO_2_ and 95% O_2_).

##### Patch-Clamp Recordings

Recordings were carried out in ACSF at a temperature of 36 °C. The intracellular recording solution was prepared with the following concentrations (in mM): 134 potassium gluconate, 6 KCl, 10 HEPES, 4 NaCl, 4 Mg_2_ATP, 0.3 NaGTP, 14 phosphocreatine di(tris), and 0.5 EGTA. Whole-cell current-clamp recordings were performed using a Dagan BVC-700A amplifier. Patch pipettes with thin-wall glass (1.5/1.0 mm OD/ID, WPI) and resistances ranging from 3 to 7 MΩ were used. Pipette capacitance was neutralized pre-break-in. Series resistance was maintained within the balanced range of 10 to 25 MΩ. Liquid junction potential was not corrected. Current signals were digitized at 20 kHz and filtered at 10 kHz (Prairie View). Gap-free recordings of central amygdala neurons expressing both GCaMP8s and ChrimsonR served as a baseline. Cells were then stimulated at 2, 10, and 20 Hz using a connectorized LED (Doric CLED_595, 595 nm wavelength - 8.5 mW with a 200 µm NA 0.53 fiber).

### QUANTIFICATION AND STATISTICAL ANALYSIS

#### Behavioral data analysis

All statistical analyses were performed in GraphPad Prism 9. Significance levels were indicated as follows: *: P<0.05, **:P<0.01, ***: P<0.001. Descriptive statistical results were presented as mean ± standard error. Mice were randomly assigned to control and experimental conditions. Data recording and analysis was performed either automatically or by an individual blind to experimental conditions. No sexual dimorphism in either histology or behavioral results was observed in the study, therefore results from males and females were grouped for analysis. For behavioral experiments, 2-tailed paired or unpaired t tests and repeated measures two-way ANOVAs were used when appropriate with poshoc tests correcting for multiple comparisons (Holm-Šídák). Welch’s correction or the Gessier-Greenhouse correction were used to account for unequal variance. Details on statistical tests and results can be found in Table S1.

#### FP data analysis

FP data was detrended to correct for photobleaching, then motion corrected by fitting the reference signal (410 nm) with the detrended activity signal (470 nm). The fluorescence change (ΔF/F) was computed by subtracting the fitted reference signal from the detrended activity signal and dividing by the fitted reference signal. Z-scored ΔF/F traces and confidence intervals were computed by bootstrap estimation. As the bi-cistronic vector generated low levels of GCaMP expression, the resulting FP signal was noisy. We computed a 95% confidence interval for each trace, and only the calcium peaks that crossed that threshold were extracted in our analysis. We also computed the area under curve (AUC) of the extracted peaks. The time points of entry into each chamber were measured from the behavioral videos. To account for the variable time spent in each chamber before and after conditioning, peaks numbers and AUC values were normalized by the duration of the stay in each chamber. Z-scored ΔF/F traces were averaged over 30s periods before and after entry into a chamber.

##### Statistics for FP

At the subject level, we employed a non-parametric approach to analyze peaks rate, duration-normalized AUC and epoch-averaged ΔF/F, given the dependency and the unknown distribution of these variables. We performed pairwise comparisons of median ranks between contexts (preferred vs placebo chamber) or conditioning (before vs after) using a bootstrap method (10000 iterations) to compute p-values, applying a Bonferroni correction to adjust the significance level for multiple comparison. At the population level, we use a multi-factor analysis of variance (ANOVA), comparing subjects, recording sides, pre/post-entry in a chamber, placebo vs preferred context, and before vs after conditioning. We applied a Bonferroni correction to adjust for multiple post-hoc comparisons. Details on statistical tests and results can be found in Tables S1 and S2.

