## Supplementary Material for "Reverse engineering placebo analgesia"

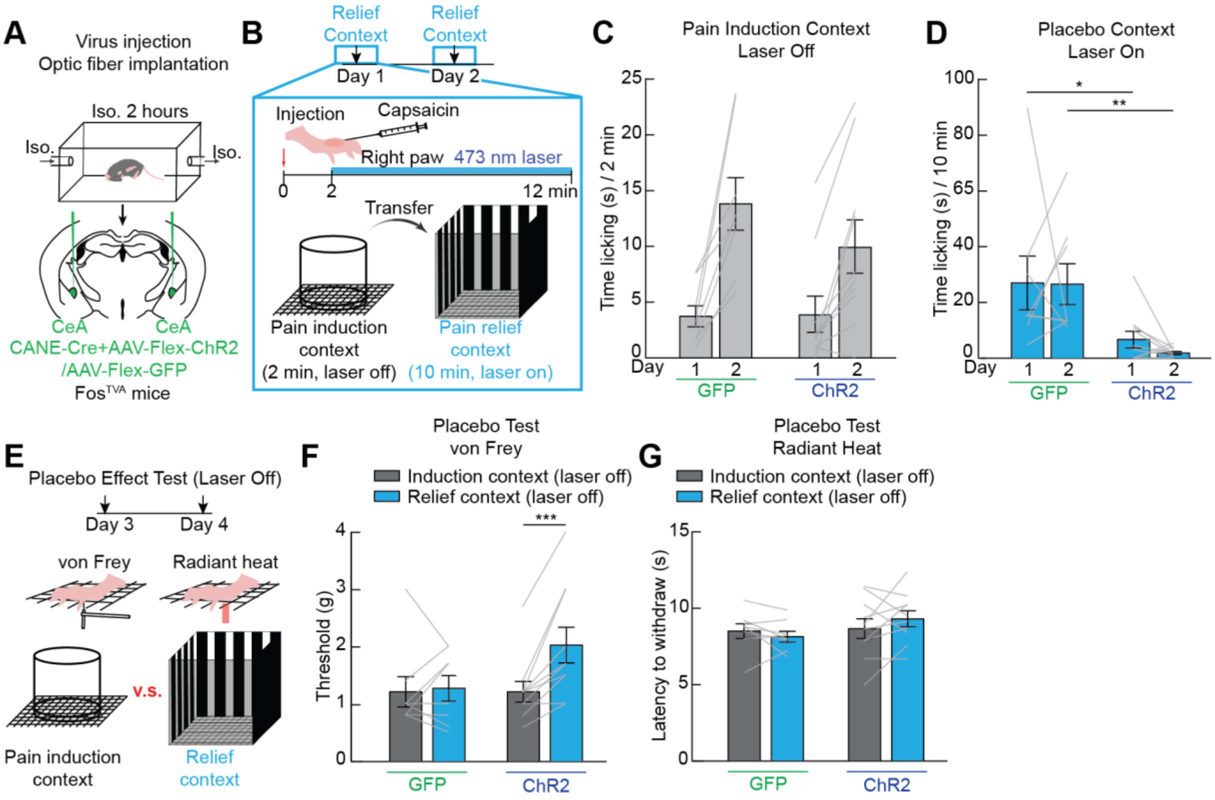


**Figure S1, related to Figure 1. Engineering placebo analgesia with a central pain suppressing circuit in a two-day acute pain model.** (**A**) Channelrhodopsin or control GFP was expressed in $CeA_{GA}$ neurons by exposing mice to isoflurane anesthesia and then injecting vectors that express in recently activated neurons using the CANE method. (**B**) Experimental timeline used to condition animals to associate a context with $CeA_{GA}$ activation-induced analgesia in an acute pain model (intraplantar capsaicin). (**C**) Time spent licking the injected paw in the induction chamber before laser stimulation (n=8-10/group, two-way repeated measures ANOVA, main effect of virus n.s.). (**D**) Time spent licking the injected paw in the conditioning chamber (n=8-10/group, two-way repeated measures ANOVA, main effect of virus p<0.001). (**E**) Experimental timeline used to test placebo analgesia by measuring mechanical (von Frey) and heat sensitivity in the induction chamber and relief context used during conditioning. (**F**) Withdrawal threshold during the von Frey test in GFP- and ChR2-expressing mice in the induction chamber (grey) and relief context (blue, n=8-10/group, repeated measures two-way ANOVA, group x context interaction p<0.05). (**G**) Latency to withdraw from radiant heat in GFP- and ChR2-expressing mice in the induction chamber (grey) and relief context (blue, n=8-10/group, repeated measures two-way ANOVA, n.s.). Data are depicted as mean ± SEM. Grey lines represent individual mice. *p<0.05, **p<0.01, ***p<0.001.

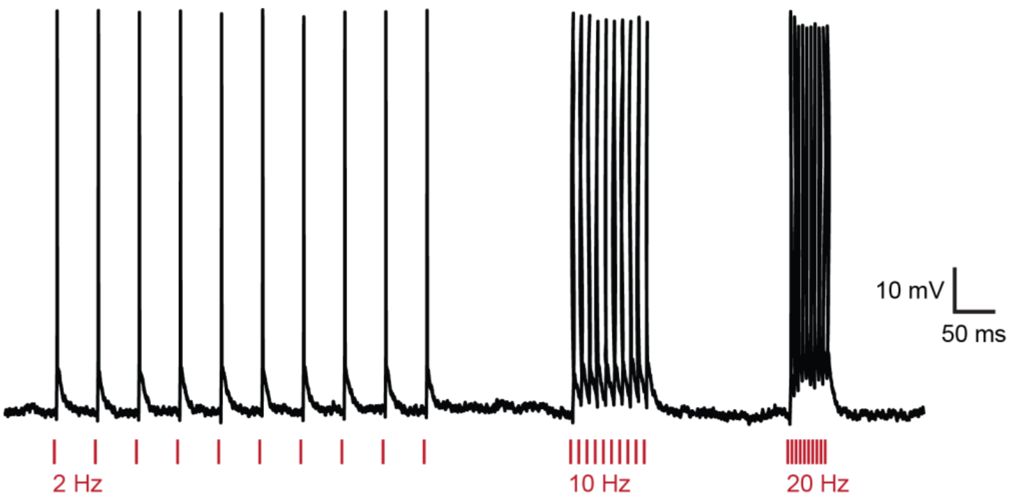


**Figure S2, related to Figure 5. Validation of ChrimsonR-mediated opto-activation of CeA neurons infected with the bicistronic AAV-DiO-jGCaMP8s-P2A-ChrimsonR virus.** ChrimsonR-expressing CeA neurons fire action potentials in response to optical stimulation at 2, 10, and 20 Hz. Scale bar: 10 mV, 50 ms.

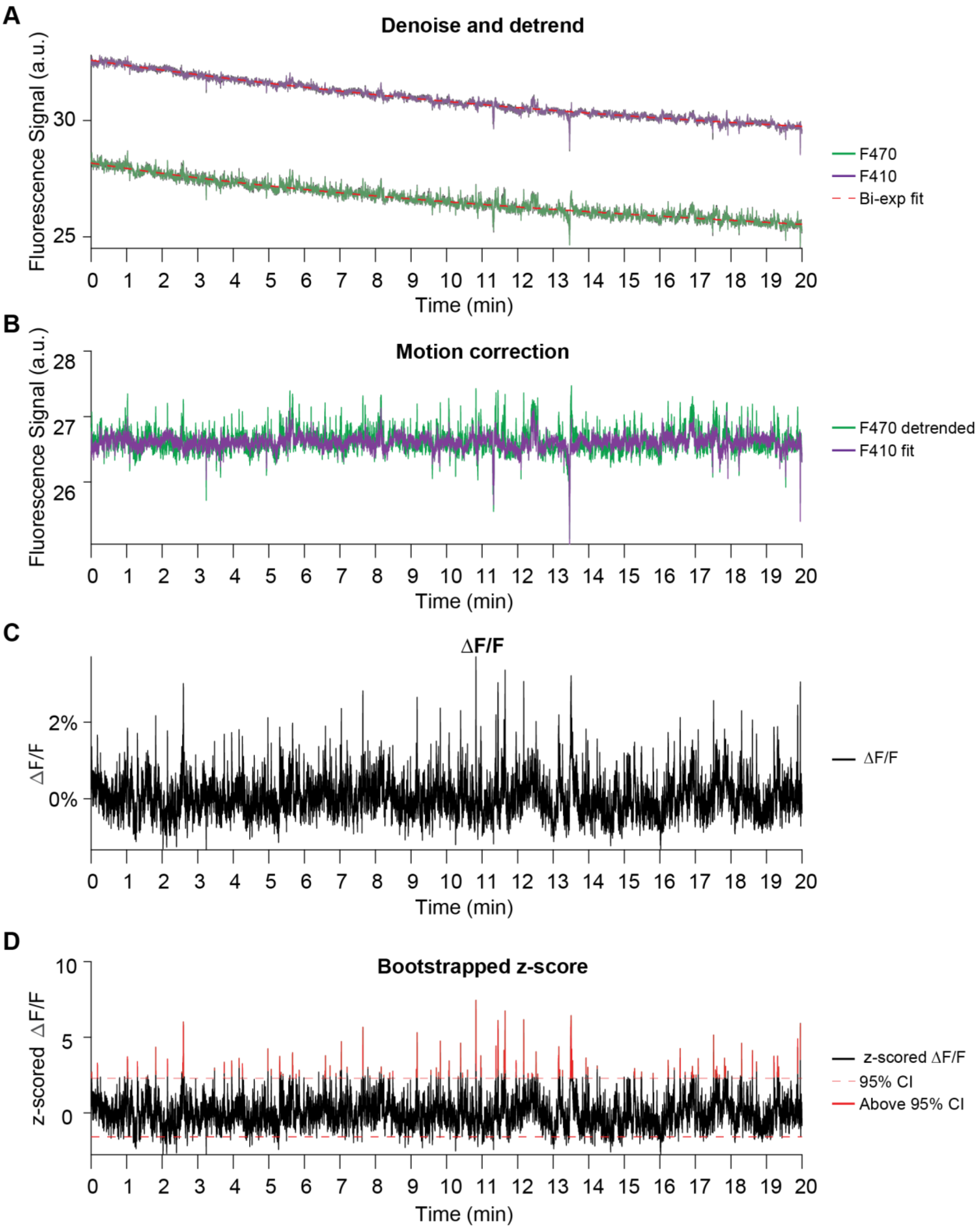
A
Denoise and detrend
Fluorescence Signal (a.u.)
30
25
0 1 2 3 4 5 6 7 8 9 10 11 12 13 14 15 16 17 18 19 20
Time (min)
F470
F410
Bi-exp fit
B
Motion correction
Fluorescence Signal (a.u.)
28
27
26
0 1 2 3 4 5 6 7 8 9 10 11 12 13 14 15 16 17 18 19 20
Time (min)
F470 detrended
F410 fit
C
ΔF/F
ΔF/F
2%
0%
0 1 2 3 4 5 6 7 8 9 10 11 12 13 14 15 16 17 18 19 20
Time (min)
ΔF/F
D
Bootstrapped z-score
z-scored ΔF/F
10
5
0
0 1 2 3 4 5 6 7 8 9 10 11 12 13 14 15 16 17 18 19 20
Time (min)
z-scored ΔF/F
95% CI
Above 95% CI

**Figure S3, related to Figure 5. Fiber photometry analysis.** (**A**) Calcium-dependent (470 nm, green) and isosbestic (410 nm, purple) jGCaMP8s signals were recorded simultaneously in $CeA_{GA}$ neurons. Denoised traces are overlayed on raw signals (in black). A bi-exponential curve (dashed red lines) was fitted to each trace and subtracted to obtain detrended signals. (**B**) Motion correction is performed by fitting the control (isosbestic, purple) channel with the calcium signal (green). (**C**) The fluorescence change (ΔF/F) is then computed by subtracting the fitted reference signal from the detrended activity signal and dividing by the fitted reference signal. (**D**) Z-scored ΔF/F traces and confidence intervals are computed by bootstrap estimation. Calcium event peaks that cross the 95% confidence threshold are considered for further analysis.

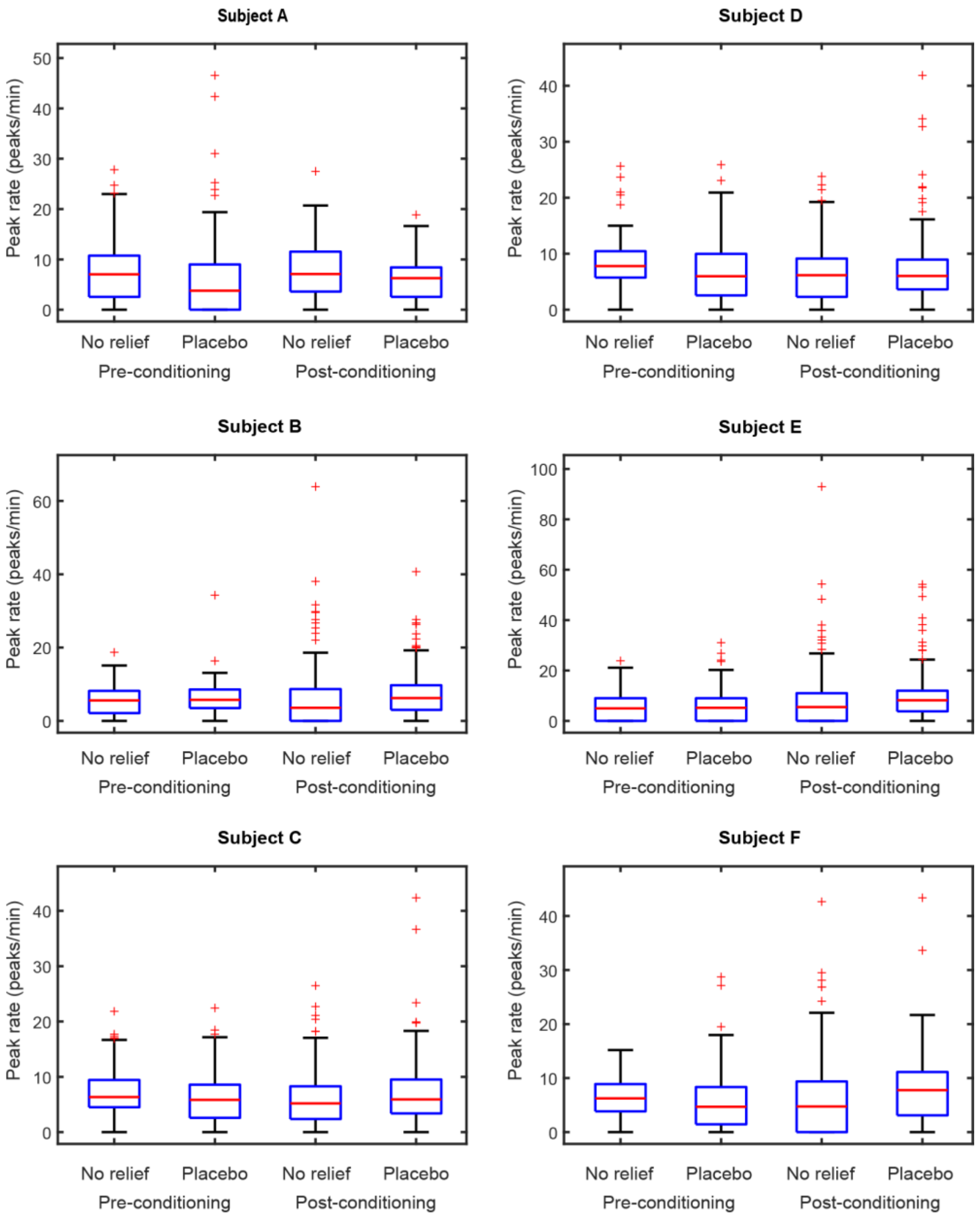


**Figure S4, related to Figure 5. Calcium event rate is unchanged after conditioning in individual mice.** Median calcium peak rate (red line in each box plot, both sides of $CeA_{GA}$ combined), for all subjects, before and after conditioning, for no-relief and placebo contexts. Bottom and top edges of boxes: 25th and 75th percentiles. Verticals whiskers encompass all data points not considered outliers (outliers are indicated as red '+' symbols). No difference between pairs of conditions was found for any individual subject (bootstrap permutation test of difference in median ranks, with Bonferroni correction for multiple comparison, n.s.).

**Table S1. Statistical tests and results.**

| Figure | Sample Size | Statistical Test | Result |
|---|---|---|---|
| 1C | N = 5-8/group | RM two-way ANOVA | |
| | | Factor 1: group | F(1,11) = 0.8783, p=0.3688 |
| | | Factor 2: time | F(3.38, 37.16) = 9.762, ***p<0.001 |
| | | Interaction: group x time | F(5,55) = 1.309, p=0.2736 |
| 1D | N = 5-8/group | RM two-way ANOVA | |
| | | Factor 1: group | F(1,11) = 3.625, p=0.0834 |
| | | Factor 2: time | F(2.88, 31.67) = 8.361, ***p<0.001 |
| | | Interaction: group x time | F(5,55) = 5.702, ***p<0.001 |
| | | Posthoc comparison: Holm-Sidak | |
| | | GFP: Day 1 vs. Day 2 | p=0.9566 |
| | | GFP: Day 3 vs. Day 4 | p=0.4796 |
| | | GFP: Day 5 vs. Day 6 | p=0.3365 |
| | | ChR2: Day 1 vs. Day 2 | *p=0.0282 |
| | | ChR2: Day 3 vs. Day 4 | p=0.0582 |
| | | ChR2: Day 5 vs. Day 6 | p=0.0593 |
| 1F | N = 5-8/group | RM two-way ANOVA | |
| | | Factor 1: group | F(1,11) = 12.25, **p=0.0050 |
| | | Factor 2: time | F(1,11) = 16.66, **p=0.0018 |
| | | Interaction: group x time | F(1,11) = 14.68, **p=0.0028 |
| | | Posthoc comparison: Holm-Sidak | |
| | | GFP: no relief vs. relief context | p = 0.9847 |
| | | ChR2: no relief vs. relief context | ***p<0.001 |
| 1G | N = 5-8/group | RM two-way ANOVA | |
| | | Factor 1: group | F(1,11) = 0.01305, p=0.9111 |
| | | Factor 2: time | F(1,11) = 9.934, **p=0.0092 |
| | | Interaction: group x time | F(1,11) = 8.724, *p=0.0131 |
| | | Posthoc comparison: Holm-Sidak | |
| | | GFP: no relief vs. relief context | p = 0.9904 |
| | | ChR2: no relief vs. relief context | ***p<0.001 |
| 1H | N = 5-8/group | RM two-way ANOVA | |
| | | Factor 1: group | F(1,11) = 0.6031, p=0.4538 |
| | | Factor 2: time | F(1,11) = 12.48, **p=0.0047 |
| | | Interaction: group x time | F(1,11) = 0.7116, p=0.4169 |
| | | Posthoc comparison: Holm-Sidak | |
| | | GFP: baseline vs. test | p = 0.1146 |
| | | ChR2: baseline vs. test | **p=0.0094 |
| 2B | N = 6 | RM one-way ANOVA | F(1.95, 9.73) = 16.31, ***p<0.001 |
| 2D | N = 6 | Paired t-test | *p = 0.0293 |
| 2E | N = 6 | Paired t-test | p = 0.2198 |
| 2F | N = 6 | Paired t-test | p=0.7661 |
| 3B (GFP) | N = 6 | RM two-way ANOVA | |
| | | Factor 1: filament | F(2.17, 10.86) = 27.64, ***p<0.001 |
| | | Factor 2: time | F(1,5) = 88.00, ***p<0.001 |
| | | Interaction: filament x time | F(1.59, 7.96) = 1.009, p=0.3876 |
| | | Posthoc comparison: Holm-Sidak | |
| | | L 0.16: Day 0 vs. Day 16 | p=0.1126 |
| | | L 0.6: Day 0 vs. Day 16 | ***p<0.001 |
| | | L 1.4: Day 0 vs. Day 16 | p=0.0880 |
| | | R 0.16: Day 0 vs. Day 16 | p=0.1126 |
| | | R 0.6: Day 0 vs. Day 16 | ***p<0.001 |
| | | R 1.4: Day 0 vs. Day 16 | p=0.0880 |
| 3B (ChR2) | N = 8 | RM two-way ANOVA | |
| | | Factor 1: filament | F(3.34, 23.37) = 22.22, ***p<0.001 |

| | | | |
|---|---|---|---|
| | | Factor 2: time<br>Interaction: filament x time<br>Posthoc comparison: Holm-Sidak<br>L 0.16: Day 0 vs. Day 16<br>L 0.6: Day 0 vs. Day 16<br>L 1.4: Day 0 vs. Day 16<br>R 0.16: Day 0 vs. Day 16<br>R 0.6: Day 0 vs. Day 16<br>R 1.4: Day 0 vs. Day 16 | F(1,7) = 21.11, **p=0.0025<br>F(3.092, 21.64) = 1.745, p=0.1866<br><br>p=0.3042<br>p=0.1641<br>p=0.3164<br>p=0.0922<br>p=0.0638<br>p=0.2712 |
| 3D (GFP) | N = 6 | RM two-way ANOVA<br>Factor 1: filament<br>Factor 2: context<br>Interaction: filament x context<br>Posthoc comparison: Holm-Sidak<br>L 0.16: no relief vs. relief context<br>L 0.6: no relief vs. relief context<br>L 1.4: no relief vs. relief context<br>R 0.16: no relief vs. relief context<br>R 0.6: no relief vs. relief context<br>R 1.4: no relief vs. relief context | <br>F(5,25) = 17.29, ***p<0.001<br>F(1,5) = 0.500, p=0.5111<br>F(5,25) = 1.547, p=0.2115<br><br>p=0.8430<br>p=0.4941<br>p=0.8224<br>p=0.7643<br>p=0.3846<br>p=0.8430 |
| 3D (ChR2) | N = 8 | RM two-way ANOVA<br>Factor 1: filament<br>Factor 2: context<br>Interaction: filament x context<br>Posthoc comparison: Holm-Sidak<br>L 0.16: no relief vs. relief context<br>L 0.6: no relief vs. relief context<br>L 1.4: no relief vs. relief context<br>R 0.16: no relief vs. relief context<br>R 0.6: no relief vs. relief context<br>R 1.4: no relief vs. relief context | <br>F(5,35) = 28.40, ***p<0.001<br>F(1,7) = 77.48, ***p<0.001<br>F(5,35) = 1.557, p=0.1979<br><br>*p=0.0154<br>***p<0.001<br>**p=0.0057<br>**p=0.0017<br>***p<0.001<br>*p=0.0278 |
| 5B | N = 6 | Paired t-test | **p=0.0092 |
| 5C | N = 6 | RM two-way ANOVA<br>Factor 1: filament<br>Factor 2: context<br>Interaction: filament x context<br>Posthoc comparison: Holm-Sidak<br>L 0.16: no relief vs. relief context<br>L 0.6: no relief vs. relief context<br>L 1.4: no relief vs. relief context<br>R 0.16: no relief vs. relief context<br>R 0.6: no relief vs. relief context<br>R 1.4: no relief vs. relief context | <br>F(3.27, 16.36) = 37.69, ***p<0.001<br>F(1,5) = 260.9, ***p<0.001<br>F(2.28, 11.43) = 1.925, p=0.1881<br><br>*p=0.0136<br>*p=0.0136<br>*p=0.0136<br>***p<0.001<br>***p<0.001<br>p=0.0778 |
| 5E | N = 6 | n-way ANOVA<br>Factor 1: subject<br>Factor 2: recording side<br>Factor 3: pre/post entry<br>Factor 4: placebo/no-relief<br>Factor 5: before/after conditioning<br>Posthoc comparison: Bonferroni<br>Placebo post-conditioning vs No-relief post-conditioning<br>Placebo post-conditioning vs Placebo pre-conditioning | <br>F(5) = 0.77, p=0.569<br>F(1) = 0.49, p=0.48<br>F(1) = 0.96, p=0.33<br>F(1) = 1.73, p=0.19<br>F(1) = 0.14, p=0.71<br><br>p=1<br>p=1 |

| | | | |
|---|---|---|---|
| | | Placebo post-conditioning vs No-relief pre-conditioning<br>No-relief post-conditioning vs Placebo pre-conditioning<br>No-relief post-conditioning vs No-relief pre-conditioning<br>Placebo pre-conditioning vs No-relief pre-conditioning | p=1<br>p=1<br>p=1<br>p=1 |
| 5F | N = 6 | n-way ANOVA<br>Factor 1: subject<br>Factor 2: recording side<br>Factor 3: placebo/no-relief<br>Factor 4: before/after conditioning<br>Posthoc comparison: Bonferroni<br>No-relief pre-conditioning vs Placebo pre-conditioning<br>No-relief pre-conditioning vs No-relief post-conditioning<br>No-relief pre-conditioning vs Placebo post-conditioning<br>Placebo pre-conditioning vs No-relief post-conditioning<br>Placebo pre-conditioning vs Placebo post-conditioning<br>No-relief post-conditioning vs Placebo post-conditioning | <br>F(5) = 16.83, ***p<0.001<br>F(1) = 5.9, p=0.015<br>F(1) = 0.39, p=0.53<br>F(1) = 0.23, p=0.63<br><br>p=1<br>p=0.79<br>p=1<br>p=1<br>p=0.18<br>p=0.06 |
| 5G | N = 6 | n-way ANOVA<br>Factor 1: subject<br>Factor 2: recording side<br>Factor 3: placebo/no-relief<br>Factor 4: before/after conditioning<br>Posthoc comparison: Bonferroni<br>No-relief pre-conditioning vs Placebo pre-conditioning<br>No-relief pre-conditioning vs No-relief post-conditioning<br>No-relief pre-conditioning vs Placebo post-conditioning<br>Placebo pre-conditioning vs No-relief post-conditioning<br>Placebo pre-conditioning vs Placebo post-conditioning<br>No-relief post-conditioning vs Placebo post-conditioning | <br>F(5) = 1.18, p=0.315<br>F(1) = 9.77, **p=0.0017<br>F(1) = 0.31, p= 0.58<br>F(1) = 4.0, p= 0.04<br><br>p=1<br>p=1<br>p=0.4<br>p=1<br>**p=0.006<br>p=0.072 |

| **Figure** | **Sample Size** | **Statistical Test** | **Result** |
|---|---|---|---|
| S1C | N = 8-10/group | RM two-way ANOVA<br>Factor 1: group<br>Factor 2: time<br>Interaction: group x time | <br>F(1,16) = 0.5764, p=0.4588<br>F(1,16) = 40.82, ***p<0.001<br>F(1,16) = 2.512, p=0.1325 |
| S1D | N = 8-10/group | RM two-way ANOVA<br>Factor 1: group<br>Factor 2: time<br>Interaction: group x time<br>Posthoc comparison: Holm-Sidak<br>Day 1: GFP vs. ChR2<br>Day 2: GFP vs. ChR2 | <br>F(1,16) = 17.26, ***p<0.001<br>F(1,16) = 0.2135, p=0.6502<br>F(1,16) = 0.1415, p=0.7117<br><br>*p=0.0156<br>**p=0.0040 |

| | | | |
|---|---|---|---|
| S1F | N = 8-10/group | RM two-way ANOVA<br>Factor 1: group<br>Factor 2: time<br>Interaction: group x time<br>Posthoc comparison: Holm-Sidak<br>GFP: no relief vs. relief context<br>ChR2: no relief vs. relief context | <br>F(1,16) = 1.321, p=0.2672<br>F(1,16) = 9.248, **p=0.0078<br>F(1,16) = 6.798, *p=0.0191<br><br>p=0.7748<br>***p<0.001 |
| S1G | N = 8-10/group | RM two-way ANOVA<br>Factor 1: group<br>Factor 2: time<br>Interaction: group x time | <br>F(1,16) = 1.009, p=0.3300<br>F(1,16) = 0.1367, p=0.7165<br>F(1,16) = 1.916, p=0.1853 |
